# Influence of amino acid substitutions in capsid proteins of coxsackievirus B5 on free chlorine and thermal inactivation

**DOI:** 10.1101/2023.11.27.568777

**Authors:** Shotaro Torii, Jérôme Gouttenoire, Kiruthika Kumar, Aleksandar Antanasijevic, Tamar Kohn

## Abstract

The sensitivity of enteroviruses to disinfectants varies among genetically similar variants and coincides with amino acid changes in capsid proteins, though the effect of individual substitutions remains unknown. Here, we employed reverse genetics to investigate how amino acid substitutions in coxsackievirus B5 (CVB5) capsid proteins affect its sensitivity to free chlorine and heat treatment. Of ten amino acid changes hypothesized to coincide with free chlorine resistance, none significantly reduced the chlorine-sensitivity, indicating a minor role of the capsid composition in chlorine sensitivity of CVB5. Conversely, we observed reduced heat sensitivity in mutants with substitutions at the C-terminal region of the viral protein 1. Cryo-electron microscopy revealed that these changes affect the assembly of intermediate viral states (particle A and E), suggesting that the mechanism for reduced heat sensitivity could be related to improved molecular packing of CVB5, resulting in greater stability and/or altered dynamics of virus uncoating during infection.

## Main

Enteroviruses are non-enveloped positive single-stranded (ss) RNA viruses and are major causative viral agents of waterborne diseases. Coxsackievirus B5 (CVB5) is a genotype of the *Enterovirus* genus, which is frequently detected in clinical and wastewater surveillance^1,2^. CVB5 is further divided into two lineages according to the phylogenetic similarities of the viral protein (VP) 1 gene^3^: genogroup A, which includes the commercially available Faulkner strain, and genogroup B. Uniquely, CVB5 demonstrates a markedly lower sensitivity to common disinfectants, such as free chlorine and heat, compared to other waterborne viruses^4–9^, making it a challenging virus to control in disinfection process.

Most studies to date have employed only CVB5 Faulkner strain, to understand CVB5 inactivation kinetics^4,6,10–12^ and to investigate its inactivation mechanism^13^ by disinfectants. It is increasingly apparent, however, that genetically diverse CVB5 environmental isolates differ in their sensitivity to free chlorine and heat^7,14^. Variant-dependent sensitivity to disinfectants has also been reported for other genotypes of enterovirus^14,15^ and other disinfectants^16^, and has been found to depend on the mechanism of action exerted by the disinfectant^7,15,17^.

Free chlorine has been found to oxidize the viral capsid^18,19^, which protects the viral genome from chemical and enzymatic damage^20,21^, and has been reported to inhibit the viral attachment function, thereby partially contributing to enterovirus inactivation^13,17,22,23^. Cysteine (Cys) and methionine (Met) react with free chlorine much faster than other amino acid residues^24,25^. Thus, the differing abundance and solvent-accessibility of Cys and Met has been suggested as the rationale for the varying sensitivities to commonly used oxidants^7,15,19,26–30^. This theory is consistent with the findings on the free chlorine susceptibility of environmental isolates of CVB5, where isolates belonging to genogroup B, which contain fewer sulfur-containing amino acids in the capsid proteins, also exhibited a lower sensitivity to free chlorine compared to isolates falling within genogroup A^28^.

In contrast to the chemical changes induced by free chlorine, thermal inactivation of enterovirus is based on structural modifications. Specifically, heat treatment can induce partial disassembly of enterovirus capsids into subunits or trigger a conformational rearrangement equivalent to viral uncoating. (i.e., transition from mature closed (F) state to altered-intermediate (A) state)^31–34^, thereby resulting in virus inactivation. This latter mechanism is suggested to be the primary driver for thermal inactivation of CVB5 at 55°C^33^. Enhanced thermotolerance can therefore be achieved by improving the stability of individual capsid building blocks (VP1-4) or strengthening the interaction network at the interfaces^35^. Consequently, enhanced capsid protein interactions were proposed to explain inter-genotype differences in the thermotolerance of enteroviruses^33^. An alternative mechanism for thermotolerance was proposed for poliovirus 1 (PV1), where amino acid substitutions in the VP1 hydrophobic pocket region stabilized the capsid against inactivation by heat by limiting premature uncoating and release of viral RNA^36^. An amino acid substitution in the pocket region, M180V at VP1, was also found after experimental evolution of CVB5 to attain thermotolerance^37^.

Despite the distinctly different inactivation mechanisms exerted by free chlorine and heat, the chlorine and heat resistance of enteroviruses have been found to coincide. Among the different enterovirus strains investigated in our past work^7^, the least chlorine-sensitive genotypes (CVB1 and CVB5) were also the least heat-sensitive, and the most chlorine-sensitive genotypes (echovirus 11; E11) were also the most heat-sensitive. Furthermore, a reduced sensitivity to free chlorine was also observed when E11 was experimentally adapted to exert greater heat tolerance^38^. Finally, the amino acid change identified in heat-adapted CVB5 (M180V in VP1) was also observed in chlorine-resistant viruses^28^. However, the genetic underpinnings of the reduced chlorine-sensitivity, heat-sensitivity, and the coincidence of the two remain unknown.

Identifying the role of individual amino acid substitutions on disinfection tolerance has been challenging, as resistant mutants typically exhibit several changes simultaneously. A powerful tool to overcome this challenge is the use of a reverse genetics system, which allows for selectively introducing individual mutations in the genome of enterovirus. While extensively used in studying viral infection and vaccine development^39–41^, reverse genetics has rarely been applied to understanding disinfection susceptibility.

Here, we employed a reverse genetic system for CVB5 and assessed how each amino acid substitution in capsid proteins affects its sensitivity to free chlorine and heat. We thereby focused on 10 amino acid substitutions identified between the two genogroups A and B that were hypothesized to reduce chlorine- and heat-sensitivity. Our findings showed that none of the substitutions, including those involving sulfur-containing residues, significantly reduced the chlorine-sensitivity of CVB5. Conversely, we observed reduced heat sensitivity in mutants with substitutions at the C-terminal region of the VP1. This information was augmented by structural analysis of a disinfection-tolerant mutant by cryo-electron microscopy (Cryo-EM), showing several local structural changes that can be linked to the introduced mutations and provide possible mechanism behind reduced heat sensitivity.

### Characterization of engineered mutants

To identify the amino acid changes between these two genogroups, the amino acid sequences in the capsid proteins (i.e., VP1-VP4) of previously tested CVB5 variants (Accession No: MW015045 - MW015056, and AF114383)^28^, seven of which belong to genogroup A and six of which belong to genogroup B, were aligned, (Table 1). In the capsid protein region, a total of thirteen conserved changes were observed between the two genogroups and six additional changes were found to be common in all except for one variant. Six amino acid substitutions are found in VP1, six in VP2, four in VP3 and three in VP4. Of these, ten were deemed of high interest. Specifically, nine substitutions, including VP1.V156I, VP1.D276E, VP1.T279A, VP2.L137I, VP2.E156D, VP2.S160T, VP2.K260R, VP3.M63L, VP3.T88I, are located on the capsid surface and are hence easily accessible to free chlorine. Among them, VP3.M63L involves the substitution of a sulfur-containing by a more chemically stable amino acid. An additional such substitution is found at a position buried within the VP1 protein, VP1.M180I. These latter two amino acid substitutions were hypothesized to be the most effective in reducing sensitivity to free chlorine.

**Table 1.**
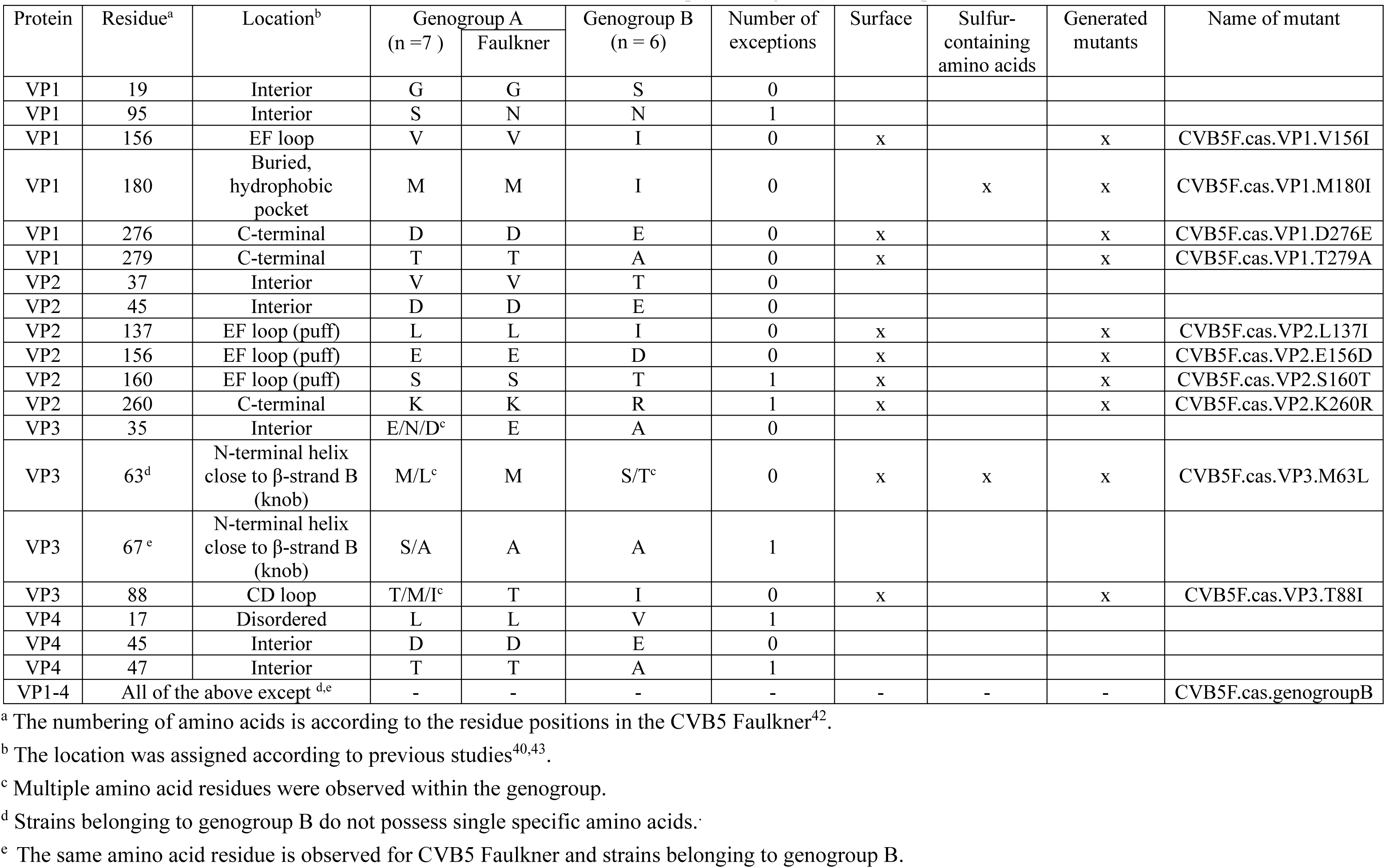
Substitutions of amino acids residues between genogroup A and B of CVB5, and description of mutants produced. CVB5F.cas corresponds to the ner strain with a *Cla*I restriction site was introduced at a non-structural protein region (nucleotide position 3340).

| Protein | Residue<sup>a</sup> | Location<sup>b</sup> | Genogroup A<br>(n = 7) |  | Genogroup B<br>(n = 6) | Number of<br>exceptions | Surface | Sulfur-containing<br>amino acids | Generated<br>mutants | Name of mutant |
| --- | --- | --- | --- | --- | --- | --- | --- | --- | --- | --- |
|  |  |  |  | Faulkner |  |  |  |  |  |  |
| VP1 | 19 | Interior | G | G | S | 0 |  |  |  |  |
| VP1 | 95 | Interior | S | N | N | 1 |  |  |  |  |
| VP1 | 156 | EF loop | V | V | I | 0 | x |  | x | CVB5F.cas.VP1.V156I |
| VP1 | 180 | Buried,<br>hydrophobic<br>pocket | M | M | I | 0 |  | x | x | CVB5F.cas.VP1.M180I |
| VP1 | 276 | C-terminal | D | D | E | 0 | x |  | x | CVB5F.cas.VP1.D276E |
| VP1 | 279 | C-terminal | T | T | A | 0 | x |  | x | CVB5F.cas.VP1.T279A |
| VP2 | 37 | Interior | V | V | T | 0 |  |  |  |  |
| VP2 | 45 | Interior | D | D | E | 0 |  |  |  |  |
| VP2 | 137 | EF loop (puff) | L | L | I | 0 | x |  | x | CVB5F.cas.VP2.L137I |
| VP2 | 156 | EF loop (puff) | E | E | D | 0 | x |  | x | CVB5F.cas.VP2.E156D |
| VP2 | 160 | EF loop (puff) | S | S | T | 1 | x |  | x | CVB5F.cas.VP2.S160T |
| VP2 | 260 | C-terminal | K | K | R | 1 | x |  | x | CVB5F.cas.VP2.K260R |
| VP3 | 35 | Interior | E/N/D<sup>c</sup> | E | A | 0 |  |  |  |  |
| VP3 | 63<sup>d</sup> | N-terminal helix<br>close to β-strand B<br>(knob) | M/L<sup>c</sup> | M | S/T<sup>c</sup> | 0 | x | x | x | CVB5F.cas.VP3.M63L |
| VP3 | 67<sup>e</sup> | N-terminal helix<br>close to β-strand B<br>(knob) | S/A | A | A | 1 | | | | |
| VP3 | 88 | CD loop | T/M/I<sup>c</sup> | T | I | 0 | x |  | x | CVB5F.cas.VP3.T88I |
| VP4 | 17 | Disordered | L | L | V | 1 |  |  |  |  |
| VP4 | 45 | Interior | D | D | E | 0 |  |  |  |  |
| VP4 | 47 | Interior | T | T | A | 1 |  |  |  |  |
| VP1-4 | All of the above except <sup>d,e</sup> |  | - | - | - | - | - | - | - | CVB5F.cas.genogroupB |

<sup>a</sup> The numbering of amino acids is according to the residue positions in the CVB5 Faulkner<sup>42</sup>.
<sup>b</sup> The location was assigned according to previous studies<sup>40,43</sup>.
<sup>c</sup> Multiple amino acid residues were observed within the genogroup.
<sup>d</sup> Strains belonging to genogroup B do not possess single specific amino acids.
<sup>e</sup> The same amino acid residue is observed for CVB5 Faulkner and strains belonging to genogroup B.

To investigate the effect of the ten selected amino acid substitutions, we engineered a total of twelve mutants. We first constructed the clone CVB5F.cas, which served as the control strain, and which differed from the Faulkner strain only by a single residue in the non-structural region (see Methods). We then constructed ten infectious cDNA clones harboring each critical amino acid substitution individually. Finally, a construct harboring all the genogroup B-specific amino acid substitutions in the capsid proteins, which the CVB5 Faulkner strain does not possess, was prepared (see Table 1). The clones are referred to as CVB5F.cas.VP*n*.X*m*Y, where an amino acid at residue number *m* in VP*n* region is replaced from X to Y, respectively. The clone containing all genogroup B-specific amino acid substitutions is termed CVB5F.cas.genogrouB. All constructs successfully produced infectious progeny viruses at comparable titers, with the final concentration of purified virus stocks ranging from 6.5 log_10_ to 7.6 log_10_ MPN mL^-1^.

### Sensitivity to free chlorine

Generated mutants were tested for chlorine-sensitivity by measuring inactivation curves in bench-scale experiments. Experimental data for free chlorine inactivation are provided in Figure S1. Estimated inactivation rate constants are shown in Figure 1. The inactivation rate constant for CVB5F.cas was 4.3 mg^-1^ min^-1^ L. The rate constants for other mutants ranged from 3.4 to 5.9 mg^-1^ min^-1^ L. Unexpectedly however, none of the mutants, including those substituting sulfur-containing amino acid and the one harboring all the genogroup B-specific amino acid substitutions in the capsid proteins, exhibited a significantly different inactivation rate constant compared to CVB5F.cas (*P* > 0.05, ANOVA). These results suggest that the introduced amino acid substitutions are not responsible for the reduced chlorine-sensitivity of CVB5 in genogroup B. The presence of sulfur-containing amino acids on the capsid’s exterior has been speculated to influence the sensitivity to free chlorine. Our findings, however, show that the decreased number of sulfur-containing amino acids in the capsid proteins does not lead to reduced chlorine-sensitivity.

**Figure 1.**
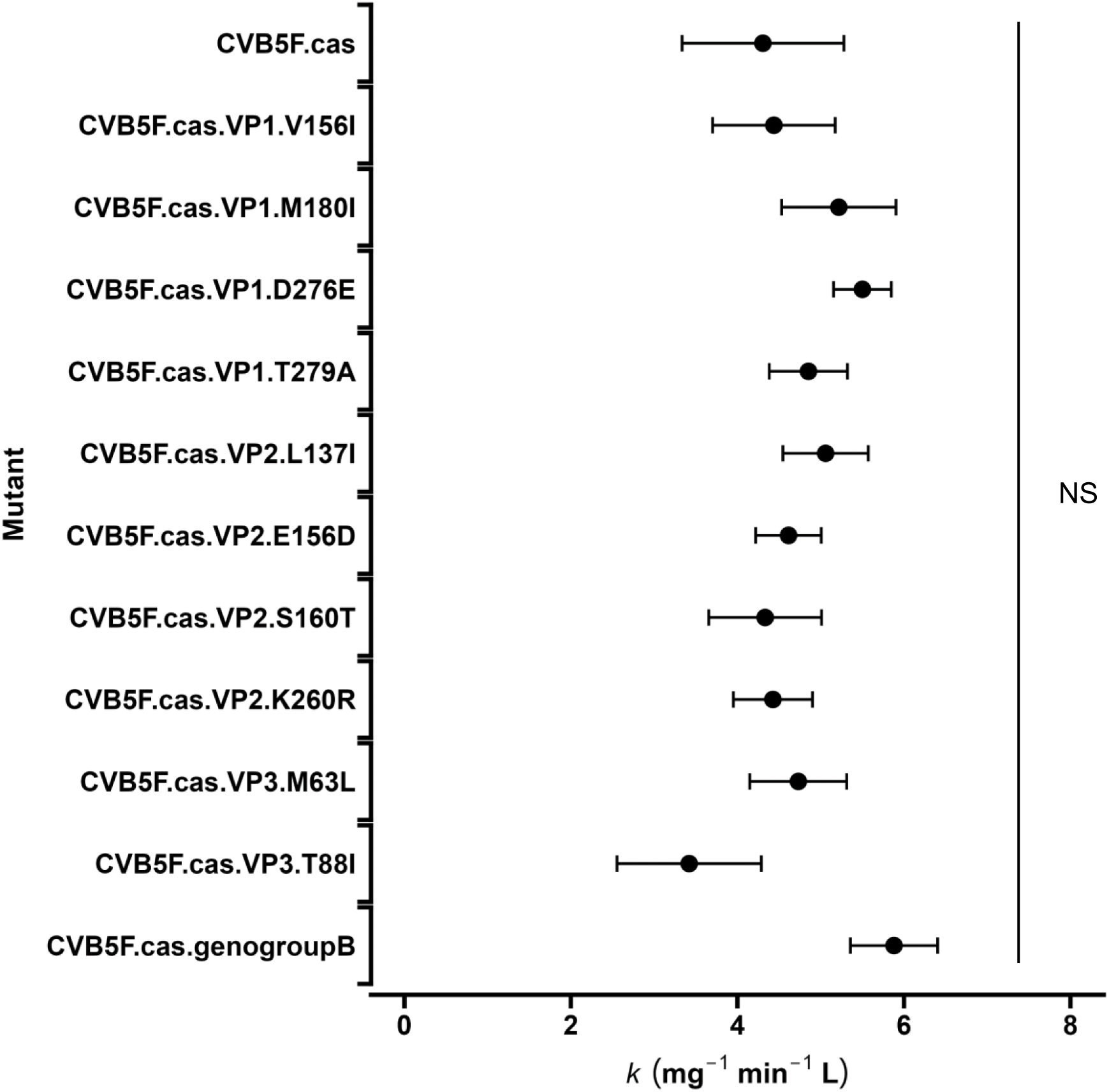
Inactivation rate constants for each mutant by free chlorine. Data are shown as mean ± 95% confidence interval on the rate constant, based on duplicate experiments. An ANCOVA analysis suggests that the rate constants are not statistically different among the mutants. NS = not significant.

### Sensitivity to heat

The mutants were tested for heat-sensitivity by exposing them to 50°C for 20 s. Experimental inactivation data of heat treatment is provided in Figure 2. The inactivation of CVB5F.cas was 1.9 ± 0.2 log_10_, with a range of 0.34 ± 0.56 to 2.1 ± 0.4 log_10_ for the other mutants. Statistically significant lower inactivation was observed for CVB5F.cas.VP1.D276E and CVB5F.cas.VP1.T279A compared to CVB5F.cas. (ANOVA with Dunnett’s post-hoc analysis; *P* < 0.05, *P* < 0.01, respectively). Consistently, the inactivation of CVB5F.genogroupB, including the two aforementioned substitutions, was also significantly lower (*P* < 0.01). Moreover, the results were consistent with previous data reporting that all the CVB5 variants containing the two substitutions exhibited lower heat sensitivity compared to CVB5F^7^. These results indicate that even a single amino acid change can significantly alter heat-sensitivity of CVB5.

**Figure 2.**
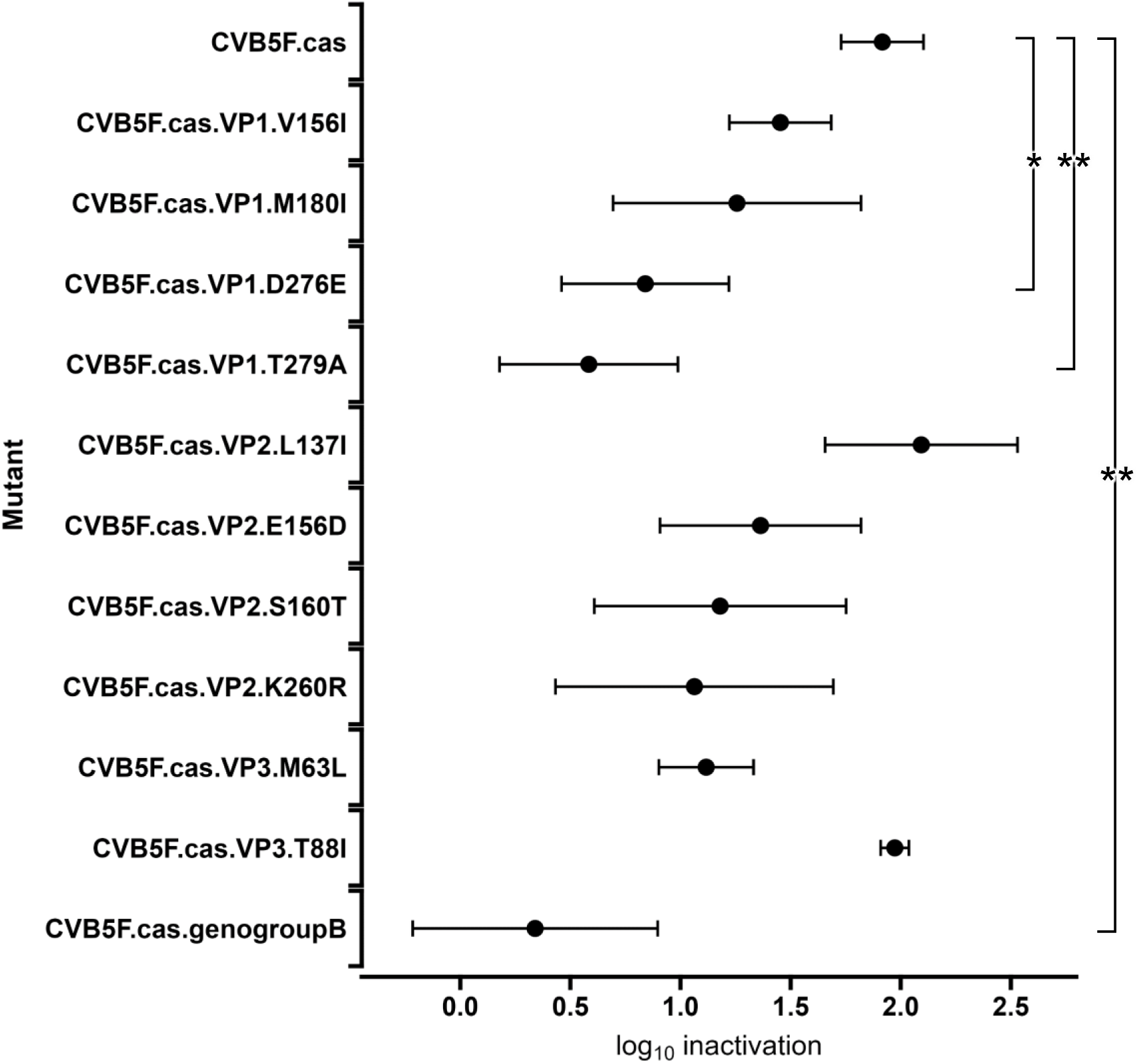
Inactivation of each mutant by heat treatment. Log inactivation is shown as the mean ± standard deviation of three replicates, with significance determined with a one-way ANOVA with Dunnett’s test for multiple comparisons (\*\**P* < 0.01, \**P* < 0.05).

To elucidate the mechanism behind reduced heat-sensitivity of the CVB5F.cas.genogroupB mutant, we subjected it to analysis by cryo-electron microscopy (cryoEM). Viruses were inactivated by formaldehyde treatment and imaged as described in the Methods section. Data collection statistics are shown in Table S1, and the data processing workflow is illustrated in Figure S2. We applied a combination of 2D and 3D classification steps in cryoSPARC package^44^ to separate the particles corresponding to mature virions (F), intermediate-altered state (A), and empty viral capsids (E). These conformational states are commonly resolved in cryoEM analyses of enteroviruses^43^, with A and E corresponding to the expanded viral particles with or without the internal viral components (non-structural proteins and genetic material). The resulting EM density maps were at 3.6Å, 2.7Å, and 2.6Å global resolution for the capsid-corresponding part of particles F, A, and E, respectively, thus allowing to build atomic models of each state (Figure 3A, Table S2).

**Figure 3.**
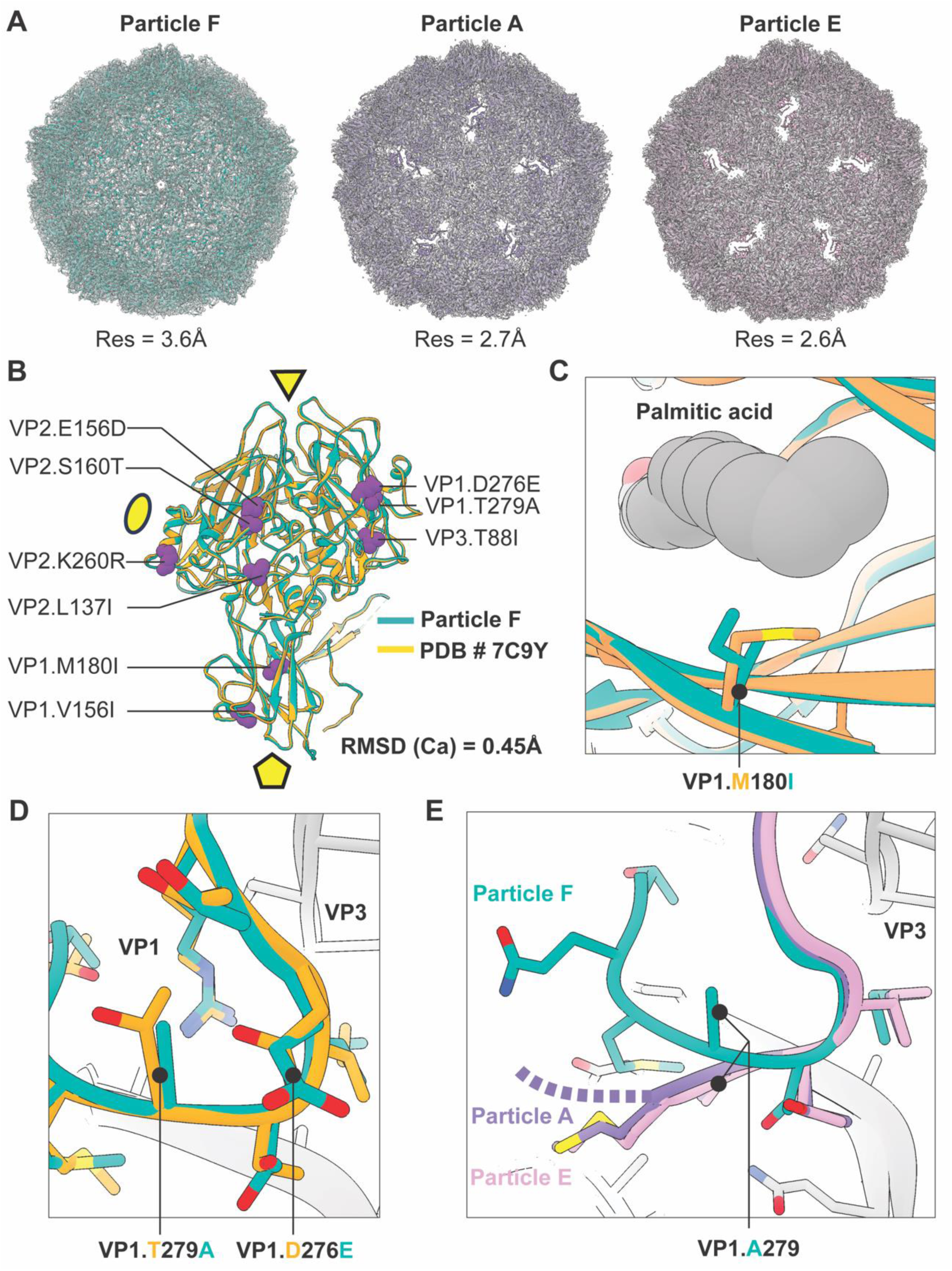
Structural characterization of CVB5F.cas.genogroupB by cryoEM. (A) Reconstructed 3D maps and models of the capsids corresponding to mature closed virion (F), intermediated-altered state (A) and empty viral capsid (E). The maps are presented as transparent gray surface and cartoon representation was used for atomic models. (B) Overlay of the CVB5F.cas.genogroupB (F) model (turquoise) and the previously published structure of the CVB5 virus (PDB ID: 7C9Y; orange) with the mutated residues presented using atomic sphere representation (magenta). For simplicity, only 1 protomer is shown with the locations of 2-, 3-, and 5-fold symmetry axis indicated with ellipse, triangle and pentamer, respectively. (C) Close-up view of the pocket in CVB5F.cas.genogroupB and non-mutated CVB5 structure with the palmitate shown in gray (sphere representation) and the alternative VP1.180 residues indicated. (D) Enlarged view of the VP1 C-terminal loop with the locations of VP1.279 and VP1.276 mutation sites indicated. Same color scheme used as in panel B. (E) Altered conformations of the structured part of VP1 C-terminus in different reconstructed states of the CVB5F.cas.genogroupB. Note the alternative conformation of the VP1.A279 residue.

Surprisingly, only ∼0.2% of the total dataset corresponded to mature virions in F state (193 particles). We believe this to be a result of formaldehyde inactivation. Nevertheless, the reconstructed atomic model closely matched the previously published CVB5 structure based on genogroup A (Figure 3B; and a previous study^43^). The root-mean-square deviation (RMSD) of Cα positions between the two structures was 0.45Å. This is consistent with the fact that most mutations were relatively conservative (L->I, E->D, S->T, K->R) and unlikely to induce major structural rearrangements. We then analyzed the locations of genogroup B mutations and if any local conformational changes arise due to their presence. None of the mutations were located at the interfaces of multiple protomers (each comprising a single copy of VP1-4). Therefore, we excluded the possibility of increased thermotolerance being due to stronger inter-protomer interactions and propose intra-protomer stabilization as a more probable mechanism.

We first focused on the VP1.M180I mutation since amino acids at this position make direct contact with the non-covalently bound palmitate residue (i.e., the pocket factor; Figure 3C). The pocket factor is essential for proper assembly of the receptor binding domain and is released during the viral entry process. While we noticed somewhat weaker density for the pocket factor compared to the previously published cryoEM map of CVB5 F-particle (EMD-30321^43^), the switch from M to I does not alter the size of the hydrophobic pocket and these two amino acids are commonly found at position 180 across the *Enterovirus* genus relevant to human infection, including 112 exemplar strains of each genotype of *Enterovirus A*, *B*, *C*, and *D* (Table S3), listed in Virus Metadata Resource^45^. This is consistent with our findings that heat-sensitivity of CVB5F.cas.VP1.M180I is not significantly different from CVB5F.cas.

The two amino acid substitutions causing the largest perturbation in relative heat-sensitivity, VP1.D276E and VP1.T279A, are located at the C-terminus of VP1. These two residues interact with each other and contribute to the external part of the VP1:VP3 interface, both through direct contacts and indirectly by stabilization of the local loop region comprising residues VP1.E272-T282 (Figure 3D). While the VP1.D276E and VP1.T279A mutations do not seem to result in any significant conformational change compared to unmutated CVB5 variant (PDB ID: 7C9Y) their pairing could affect local molecular packing and influence the dynamics of viral unfolding during infection or thermal inactivation. Consistently, this region in VP1 undergoes partial restructuring in intermediate-altered (A) and empty particle (E) states (Figure 3E), with alanine at VP1.279 facing towards the hydrophobic pocket in VP3 assembled by residues VP3.P86, VP3.A141 and VP3.Y189. Although, the C-terminus of VP1 is flexible in different enterovirus structures, the resulting conformation in A and E particles of CVB5F.cas.genogroupB is distinct from the previously published reconstruction of CVB5-A and E particles having the original VP1.D276/VP1.T279 pairing (Figure S3 and the previous study^43^), supporting that VP1.D276E and VP1.T279A mutations may affect the assembly and local dynamics of the VP1:VP3 interface.

Several other areas in particle A and E states of CVB5F.cas.genogroupB also exhibit different conformations and/or greater flexibility compared to corresponding CVB5 structures from the PDB (Figure S3). Most prominent changes were in the N-terminal region of VP1 (VP1.Q49-S60), VP2 (VP2.L42-Q52) and VP3 (I168-V181). While these discrepancies could indicate altered energy landscape of intermediate viral conformations in CVB5F.cas.genogroupB, none of the listed residue ranges contain genogroupB mutations, or create direct contacts with them. Therefore, the observed effects are either indirectly influenced by amino-acid substitutions, or potentially a consequence of formaldehyde treatment.

Altogether, based on structural analysis of the CVB5F.cas.genogroupB we propose that mutations conferring greater tolerance to thermal inactivation act by perturbing local molecular interactions which may lead to improved stability of infectious virions (F) and altered conformations of uncoating intermediates (e.g., particles A and E).

## Discussion

The impact of mutations on disinfection sensitivities has been a subject of debate, yet the direct effect of mutations has been rarely evaluated. Notable exceptions include an investigation of the sensitivity of murine norovirus to calcium hydroxide^46^, as well as the sensitivity of PV1 to chlorine^36^. Given that CVB5 is among the most resistant viruses to common disinfectants, our study employed a reverse genetics system to assess how single amino acid substitutions influence the sensitivity of this virus to free chlorine and heat, yielding several important insights.

Our free chlorine disinfection experiments showed a surprising result; the substitution of sulfur-containing amino acids by aliphatic ones did not coincide with a lowered sensitivity to free chlorine. This finding contradicts hypotheses raised by prior studies on the role of oxidizable residues in free chlorine disinfection^7,26,28,47,48^, but is consistent with reports on the absence of chlorine resistance in PV1 following a methionine to valine substitution in VP1^36^. The sensitivity to other oxidants, such as peracetic acid are also believed to be governed by the structure and protein compositions of viruses^27,29^. Our result with free chlorine implies a need to revisit this hypothesis.

We then sought alternative explanations for the genogroup-dependent CVB5 sensitivity to free chlorine. Past studies showed that another oxidant, chlorine dioxide, primarily damages the viral genome region spanning approximately from nucleotide 1 to 120, within 5’ untranslated region (5’UTR), leading to inactivation of PV1 and enterovirus 71^49^. It was also reported that the 5’UTR of hepatitis A virus is the most degraded by free chlorine across the whole genome^50^. The 5’ UTR contains a cloverleaf structure directing viral RNA replication and an internal ribosome entry site that initiates translation^51^. This suggests that the region is also essential for enterovirus infectivity.

Interestingly, a 5’UTR-based classification of the CVB5 isolates used in our previous work^28^ segregated the variants into the same clusters as the VP1-based classification, with the sole exception of CVB5 Faulkner. In contrast, when other genome regions were used as a classification basis, the clustering diverged (Table S4 and Figure S4). Moreover, the inactivation rate constants of the variants that we previously tested are also significantly different between the two 5’UTR-based genogroups (Wilcoxon-rank sum test: *P* < 0.01). Given that the genome damage induced by free chlorine also contributes to viral inactivation^13,52^, it is plausible that the mutations in 5’UTR alter the composition and the secondary structures, thereby changing chlorine-sensitivity. A total of 12 common mutations were observed between the two 5’UTR-based genogroups. Further studies should explore the impact of each mutation on the chlorine-sensitivity. On the other hand, our study found that the heat-sensitivity of CVB5 is lowered with the amino acid substitutions in the C-terminal region of VP1. This highlights the importance of characterizing the thermostability of a given viral genotype or species based on multiple variants, rather than a single strain.

Although unveiling the exact mechanism of the reduced heat-sensitivity needs further examination of uncoating mechanism of CVB5, we speculate the contribution of the stabilized VP1:VP3 interface by the substitution of VP1.D276E and VP1.T279A. A past structural analysis of coxsackievirus A7 showed that the uncoating triggered by heat treatment at 56 °C results in the rotation of VP1, which causes major conformational changes at the interfaces of the capsid proteins VP1, VP2, and VP3^53^. The substitution of VP1.D276E and VP1.T279A may thus minimize the dissociation of the VP1:VP3 interface by heat treatment, resulting in the reduced heat-sensitivity. Interestingly, an alignment analysis of 112 exemplar strains of the *Enterovirus* genus, representing all genotypes relevant to human infection, revealed no occurrence of the combination of VP1.E276 and VP1.A279 observed in CVB5 genogroup B, whereas the combination of VP1.D276 and VP1.T279 observed in CVB5 genogroup A is also observed for 15 other *Enterovirus* genotypes (e, g, PV1, CVA9, CVA1). Therefore, the reduced heat-sensitivity by this combination of amino acid substitution is assumed to be specific to CVB5 genogroup B. It is worth investigating the relationship between the amino acid substitution at C-terminal VP1 region, its effect on VP1:VP3 interface, and heat-sensitivity of viruses in further studies.

This study also suggests that the chlorine- and thermotolerance do not necessarily correlate, at least if thermotolerance is induced by amino acid substitutions in the capsid proteins. This is consistent with previous studies that found no effect of thermotolerance-inducing substitutions in the capsid protein of PV1 on chlorine resistance^36^. Instead, the two effects appear to be induced by separate sets of amino acid substitutions that co-occur in CVB5 isolates belonging to genogroup B. Nevertheless, a correlation between chlorine- and thermotolerance cannot be excluded for mutations occurring in other regions of the genome rather than capsid proteins region. In particular, future work should focus on the role of mutations in the 5’ UTR on both chlorine and heat treatment.

Finally, the employed reconstituted infectious cDNA clone features a cassette vector system where capsid protein regions can be substituted with that from a different strain, genogroup, or genotype^54^. Testing these viruses allows for investigating the fundamental cause of variant-, genogroup-, and genotype-dependent disinfection sensitivity of enterovirus. The use of reverse genetics presents significant potential for advancing our understanding of the differing sensitivity of viruses to disinfectants and their respective inactivation mechanisms.

## Methods

### Cells

Buffalo green monkey kidney (BGMK) cells were kindly provided by the Spiez laboratory (Switzerland). The cells were grown in Eagles’s Minimum Essential Medium (MEM) supplemented with 10% of fetal bovine serum (FBS; Gibco) and 1% of penicillin−streptomycin (P/S; Gibco), and were maintained in MEM supplemented with 2% FBS and 1% of P/S at 37 °C in humidified 5% CO_2_-saturated conditions.

### Construction of Infectious cDNA clone

Plasmids containing partial CVB5 Faulkner (CVB5F) genome (Accession Number: AF114383) in a pUC57 vector were purchased (GenScript) and were cloned by Gibson Assembly^55^ using Gibson Assembly Mastermix (NEB) to produce an infectious, full-length cDNA clone of CVB5F. In this CVB5F clone (later termed CVB5F.cas), a *Cla*I restriction site (Figure 4) was introduced at a non-structural protein region (nucleotide position 3340) to generate a cassette vector^40^. While this modification resulted in one amino acid change in the 2A protein (valine to leucine at amino acid position 17), the introduced substitution was accepted, as leucine 17 is present in the 2A protein of other enteroviruses, including E30 and E21^40^.

**Figure 4.**
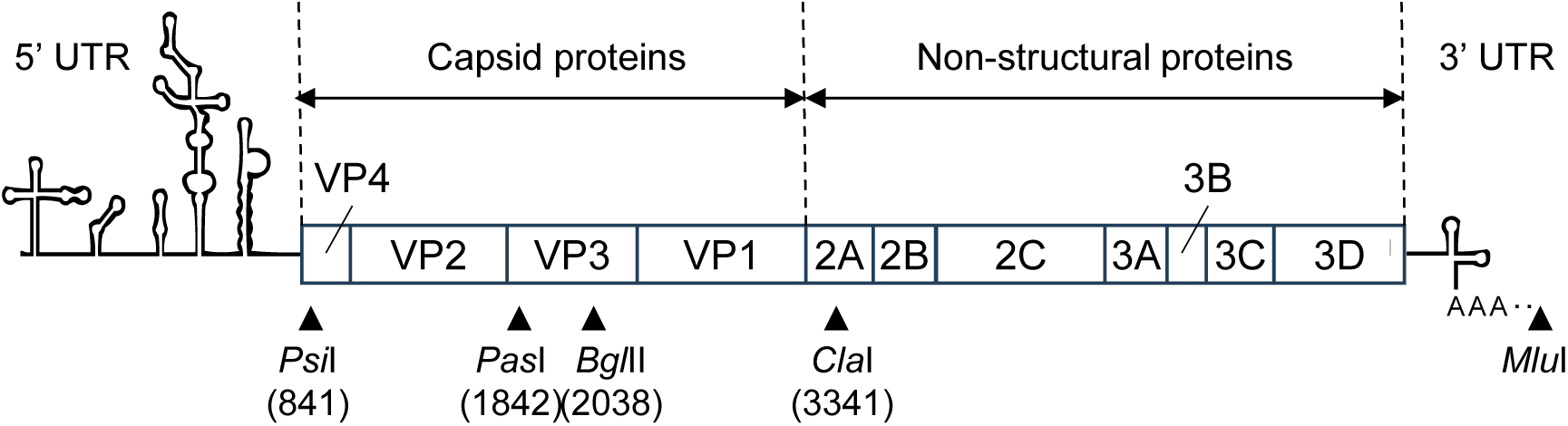
Genomic structure of CVB5F.cas. An illustration of the genome organization is shown along with the position numbers of relevant restriction sites used to construct infectious cDNA clones in this study. The position number is based on CVB5 Faulkner (accession No. AF114383).

### Mutagenesis

In addition to the CVB5F.cas clone, a total of 11 mutants were generated in this study (Table 1). Ten of them were the mutants containing single amino acid substitution and are referred to as CVB5F.cas.VP*n*.X*m*Y, where an amino acid at residue number *m* in VP*n* region is replaced from X to Y, respectively. The other simultaneously contains all the unique amino acid substitutions from genogroup A to genogroup B (i.e., all the listed substitutions except for VP3.M63L in Table 1), referred to as CVB5F.cas.genogroupB. For generating CVB5F.cas.VP*n*.X*m*Y, pCVB5F.cas was site-directed mutagenized by PCR using the primers listed in Table S5. A pair of either *Psi*I, *Pas*I, *Bgl*II, and *Cla*I restriction sites was used to linearize pCVB5F.cas (Figure 4). The PCR-amplified region containing the desired mutation was cloned by Gibson Assembly to produce each infectious cDNA clone. For pCVB5F.cas.genogroupB, pTwist Amp High Copy plasmid containing the genomic sequence of pCVB5F.cas.genogroupB flanked by *Psi*I and *Cla*I was purchased (Twist Bioscience). The region flanked by the two restriction sites were amplified by PCR and cloned into pCVB5F.cas by Gibson Assembly. All the produced constructs were propagated in competent cells (5-alpha Competent *E.coli*; NEB) and isolated by NucleoSpin Plasmid (TaKaRa). The introduced mutation was verified by Sanger sequencing.

### Generation of viruses from infectious cDNA clones

A full-length CVB5 RNA was produced by *in vitro* transcription of an *MluI*-linearized plasmid using the T7 RiboMAX™ Express Large Scale RNA Production System (Promega). Then, 0.4 – 2.6 μg of the transcribed RNA was transfected into near confluent BGMK cells prepared in T25 flask using Lipofectamine MessengerMax transfection reagent (Thermo Scientific)^23^. After incubation at 37 °C in humidified 5% CO_2_-saturated conditions for three days, the transfected cells were frozen and thawed once and harvested. The cell suspension was centrifuged at 3500 g for 15 min to pellet down cells debris. The supernatant was filtered through a 0.45 μm low protein binding durapore membrane (Merck Millipore Ltd.), and the progeny virus stock was aliquoted and stored at −20°C until use.

A 100 μL aliquot of the progeny virus was passaged in a T150 flask with confluent BGMK cells for a large-scale virus production. The passaged viral stock was recovered as described above. Then, 30 mL of the passaged virus stock was purified by sucrose-cushion ultracentrifugation followed by 0.22 μm membrane filtration as described previously^22^. The purified viral stock was stored at 4°C before use. Sanger sequence confirmed that the introduced mutation was preserved even after cell passage, except for CVB5F.cas.VP1.V156I. For this mutant, an ambiguous nucleotide, denoted Y (i.e., C or T), was observed at position 2914, but the mutation was synonymous, ensuring that the intended amino acid sequence was maintained (Table S6).

### Enumeration of infectious viruses

Infectious virus concentrations were enumerated by end point dilution assay using near confluent BGMK cells maintained in 96-well plates as described elsewhere^9^ and were quantified according to the most probable number (MPN) method^56^ using the R package {MPN}^57^. The lower limit of detection was 12 MPN mL^-1^ of sample.

### Disinfection Experiments

#### Free chlorine

The free chlorine disinfection experiment was conducted in a glass beaker in duplicate for each mutant. All experiments were conducted at a temperature-controlled room at 20°C. A free chlorine working solution was prepared by diluting sodium hypochlorite (Reactolab SA, Switzerland) in 1 mM phosphate buffer (pH 7.0). The final free chlorine concentration in the working solution ranged from 0.53 to 0.62 mg L^−1^. The free chlorine concentration was measured by the *N*, *N*-diethyl-*p*-phenylenediamine (DPD) method^58^ using a DR300 Chlorine Pocket Colorimeter (Hach Company, USA). Before each run, glass beakers were soaked with >50 mg L^−1^ of sodium hypochlorite overnight to quench the residual chlorine demand. The beakers were rinsed twice with MilliQ water and once with the chlorine working solution. Then, 50 μL of virus stock solution were spiked into 11.5 mL of the working solution under constant stirring, to achieve a starting concentration of 4.1 – 5.2 log_10_ MPN mL^-1^. A 500 μL aliquot was collected every 30 s or 45 s and mixed with 5 μL of 5,000 mg L^−1^ sodium thiosulfate (Sigma-Aldrich, Germany) to instantly quench the residual free chlorine. A total of three time-series samples plus an untreated sample (i.e., sample at time zero), were taken. After the experiment, untreated and disinfected samples were stored at 4℃ for a maximum of 24 h. The free chlorine concentration in the beaker was measured at the beginning and ten seconds after the collection of the last time-series sample. The decay in free chlorine concentration was less than 16% throughout each run. The chlorine exposure (CT value; concentration of free chlorine multiplied by contact time) for each sample was determined by integration of the time-dependent disinfectant concentration over exposure time, assuming first-order decay in free chlorine concentration between the two time points. The inactivation rate constants (*k*) (mg^-1^ min^-1^ L) were determined based on the pooled data from all replicates as the slope of −ln(*N/N_0_*) versus CT value by linear least-squares regression, where *N* is the infectious virus concentration at time T (MPN mL^−1^) and *N_0_* is the infectious virus concentration at time 0 (MPN mL^−^1)

#### Heat

Heat treatment was conducted in a thermal cycler (GeneAmp PCR system 9700, Applied Biosystems, USA) in triplicate. Five microliters of purified virus stock were spiked into thin-wall PCR tubes containing 45 μL of 1 mM phosphate buffer pre-heated at 50°C and incubated for 20 s. The incubated tubes were immediately cooled down by placing them on crushed ice, and samples were stored at 4℃ as specified above until enumeration. Log_10_ inactivation was given by −log_10_(*N/N_0_*).

### CryoEM and single particle 3D construction

#### Preparation of the virus sample for cryoEM imaging

A 160 mL aliquot of the passaged virus stock of CVB5F.cas.genogroupB was placed on 20% sucrose cushion and ultracentrifuged at 150,000 g for 3 hrs. After the supernatant was decanted, the pellet was resuspended with 500 μL of filtered phosphate-buffered saline (PBS;10 mM phosphate, 140 mM NaCl, 2.68 mM KCl, pH 7.4, Gibco). The suspension was again centrifuged at 10,000 g for 3 min to remove the carry-over debris. The supernatant was collected and amended with paraformaldehyde at 100 μg mL^-1^ and incubated at 4°C for five days to fix the samples. The fixed samples were washed with PBS using an Amicon Ultra centrifugal unit (MWCO: 100 kDa, Merck Millipore) and subjected to size exclusion chromatography to further purify the viral fraction as described elsewhere^59^. Size-exclusion chromatography was performed using a HiPrep 16/60 Sephacryl S-500 HR column (Cytiva) running in tris-based buffer (25 mM Tris-HCl, 150 mM NaCl, pH 7.5). Fractions corresponding to CVB5 were combined and concentrated to 4.5 mg ml^-1^ using Amicon Ultra centrifugal filter units with 100 kDa MWCO (Merck Millipore).

#### Grid preparation and imaging

Grids were prepared as described before^59^. Briefly, 3 µL of the purified CVB5 mutant virus sample at 4.5 mg mL^-1^ concentration was loaded onto Quantifoil R 1.2/1.3 grids (EMS), that were previously glow-discharged for 30 s in a GloCube Plus device (Quorum Technologies). Grid vitrification was performed on a Vitrobot Mark IV with the following settings: Temperature = 10 °C; Humidity = 100%; Blotting force = 0; Wait time = 10 s; Blotting time varied in the 4-6 s range. Following the blotting step, the grids were plunge-frozen into liquid ethane, cooled by liquid nitrogen. Samples were imaged on a Glacios electron microscope (Thermo Fisher Scientific) equipped with an X-FEG electron source and operating at 200 kV voltage. Images were collected with a Falcon IVi camera in the electron-event-representation (EER) format. Nominal microscope magnification was set to 150,000 X resulting in a pixel size was 0.926 Å (at the specimen plane). Automated data collection was performed using the EPU software (Thermo Fisher Scientific). Data collection information is provided in Table S1.

#### Data processing and model reconstruction

All data processing steps were performed in cryoSPARC software package^44^. Raw micrograph frames were aligned and dose-weighted using Patch Motion Correction while the CTF parameters were estimated using CTFFind^60^. Particle picking was done using a combination of blob and template picker in cryoSPARC, resulting in 161’820 total extracted particles that were then subjected to 2D classification. Bad particle picks were removed, and virus-resembling particle classes were divided into empty and full, based on the absence/presence of internal viral components in the 2D classes. “Empty” class comprised 75’212 particles (E). Particles containing internal viral components (∼28’000) were subjected to heterogeneous refinement with icosahedral symmetry imposed. Initial 3D model was generated by Ab initio reconstruction of the full 2D-cleaned dataset. Heterogeneous refinement resulted in 2 distinct subsets corresponding to the closed native virus state (F, 193 particles) and the intermediate-altered conformation (A, 27’505 particles). The F, A and E subsets were then subjected to non-uniform refinement in cryoSPARC with icosahedral symmetry imposed, and refinement of global and local CTF parameters. The resulting maps were used for the relaxation of atomic models. The entire data processing workflow is shown in Figure S3 Model building and refinement was completed using a combination of manual steps in Coot^61^ and automated steps in Rosetta^62^. Only 1 asymmetric unit was built per viral particle with icosahedral symmetry restraints imposed. Throughout the EM map we observed additional densities surrounding the side chains of histidine and cysteine residues, that cannot be assigned to any peptidic or posttranslational elements (Figure S3). We believe these to be the results of formaldehyde treatment^63^, but we did not try to approximate them with atomic models. Model validation was performed in Phenix^64^ using the MolProbity^65^ and EMRinger^66^ metrics. The resulting models and maps were deposited to the Protein Data Bank (PDB) and Electron Microscopy Data Bank (EMDB), respectively. Model statistics and PDB/EMDB accession numbers are shown in Table S2.

#### Alignment of amino acid sequences

A total of 112 exemplar virus isolates, each of which belong to a different genotype of human-infecting Enterovirus (i.e, *Enterovirus A*, *B*, *C*, and *D*) listed in Table S6, were identified from Virus Metadata Resource from International Committee on Taxonomy of Viruses^45^. All the full-length amino acid sequences of polyproteins were aligned in Geneious Prime 2023.1.2 using MAFFT plugin^67^ with default settings. The alignments were examined for the amino acid variations at position 180, 276, and 279 in VP1 region.

#### Statistical analyses

All statistical analyses were performed using R version 4.3.0^68^. Linear least-squares regression was performed with lm function to estimate inactivation rate constants. An analysis of covariance (ANCOVA) was performed with an R package {car}^69^ to compare the inactivation rate constants by free chlorine among the mutants. A one-way analysis of variance (ANOVA) with Dunnett’s post-hoc analysis was conducted with an R package {multcomp}^70^ to compare the level of inactivation after 20 s of heat treatment among the mutants. Graphs were generated with the packages {ggplot2}^71^ and {ggprism}^72^.

## Acknowledgement

This work was funded in part by the Japan Society for the Promotion of Science (JSPS) Overseas Challenge Program for Young Researchers, the Young Researchers Exchange Programme between Japan and Switzerland (EGJP_04-042020), JSPS Overseas Research Fellowships to S.T., and by EPFL discretionary funds. The work described in this paper was also supported by amfAR, The Foundation for AIDS Research, grant # 10413-73-RKVA awarded to A.A. Electron microscopy grids were vitrified using the equipment at the Protein Production and Structure Core (PTPSP) at EPFL. Electron microscopy data was collected at the Dubochet Center for Imaging (DCI) in Lausanne with assistance from Alexander Myasnikov, Emiko Uchikawa, Bertrand Beckert and Sergey Nazarov. Electron microscopy data was processed using the computational infrastructure provided by the IT department of the School of Life Sciences (SV-IT). The authors express sincere gratitude to the PTPSP, DCI and SV-IT personnel for their contribution.

## Supporting Information

The Supporting Information is available.

## Data availability

All data discussed in this manuscript are accessible via https://doi.org/10.5281/zenodo.10141286. 3D maps and models from the electron microscopy experiments have been deposited to the Electron Microscopy Databank (http://www.emdatabank.org/) and the Protein Data Bank (http://www.rcsb.org/), respectively. The accession numbers are listed in the Methods section and Table S2.

## Notes

https://zenodo.org/records/10141286

### Competing Interest Statement

The authors have declared no competing interest.

