## Supporting Information for "Influence of amino acid substitutions in capsid proteins of coxsackievirus B5 on free chlorine and thermal inactivation"

Contents (15 pages, six Tables, and four Figures)

Table S1 CryoEM data collection information

| | **CVB5F.cas.genogroupB** |
|---|---|
| **Microscope** | TFS Glacios (X-FEG) |
| **Voltage (kV)** | 200 |
| **Detector** | Falcon IVi |
| **Recording mode** | EER |
| **Magnification** | 150,000 X |
| **Movie micrograph pixel size** | 0.926 |
| **Dose rate ($e^{-}$/Å$^2$/s)** | 9.0 |
| **Total Exposure Dose ($e^{-}$/Å$^2$)** | 40 |
| **Number of EER fractions** | 40 |
| **Movie micrograph exposure time (s)** | 9.5 |
| **Total dose ($e^{-}$/Å$^2$)** | 44.7 |
| **Nominal under focus range (µm)** | 0.8 – 1.0 |
| **Number of movie micrographs** | 6653 |

Table S2 Model building and refinement information

| | F-particle | A-particle | E-particle |
|---|---|---|---|
| **EMDB ID** | EMD-18942 | EMD-18943 | EMD-18944 |
| **Map Resolution (Å)** | 3.6 | 2.7 | 2.6 |
| **Map Symmetry** | I | I | I |
| **PDB ID** | 8R5X | 8R5Y | 8R5Z |
| **Residues** | 819 | 658 | 658 |
| **Amino-acids** | 818 | 658 | 658 |
| **Ligands** | 1 | 0 | 0 |
| **RMSD Bonds** | 0.021 | 0.020 | 0.019 |
| **RMSD Angles** | 1.741 | 1.685 | 1.718 |
| **Ramachandran** | | | |
| **Outliers (%)** | 0.00 | 0.00 | 0.00 |
| **Allowed (%)** | 1.11 | 0.78 | 1.09 |
| **Favored (%)** | 98.89 | 99.22 | 98.91 |
| **Rotamer outliers** | 0.00 | 0.00 | 0.00 |
| **Clash score** | 1.20 | 1.98 | 1.29 |
| **Molprobity score** | 0.84 | 0.96 | 0.85 |
| **EMRinger score** | 3.09 | 5.22 | 4.83 |

Table S3 List of exemplar virus isolates extracted from Virus Metadata Resource from International Committee on Taxonomy of Viruses[1].

| Species | Genotype(s) | Virus name abbreviation(s) | Virus isolate designation | Virus GENBANK accession |
|---|---|---|---|---|
| *Enterovirus A* | coxsackievirus A2 | CVA2 | Fleetwood (Delaware/47) | AY421760 |
| *Enterovirus A* | coxsackievirus A3 | CVA3 | Olson (New York/48) | AY421761 |
| *Enterovirus A* | coxsackievirus A4 | CVA4 | High Point (North Carolina/50) | AY421762 |
| *Enterovirus A* | coxsackievirus A5 | CVA5 | Swartz (New York/50) | AY421763 |
| *Enterovirus A* | coxsackievirus A6 | CVA6 | Gdula | AY421764 |
| *Enterovirus A* | coxsackievirus A7 | CVA7 | Parker | AY421765 |
| *Enterovirus A* | coxsackievirus A8 | CVA8 | Donovan (New York/49) | AY421766 |
| *Enterovirus A* | coxsackievirus A10 | CVA10 | Kowalik (New York/50) | AY421767 |
| *Enterovirus A* | coxsackievirus A12 | CVA12 | Texas 12 (Texas/48) | AY421768 |
| *Enterovirus A* | coxsackievirus A14 | CVA14 | G-14 (South Africa/50) | AY421769 |
| *Enterovirus A* | coxsackievirus A16 | CVA16 | G-10 (South Africa/51) | U05876 |
| *Enterovirus A* | enterovirus A71 | EV-A71 | BrCr | U22521 |
| *Enterovirus A* | enterovirus A76 | EV-A76 | FRA91-10369 | AY697458 |
| *Enterovirus A* | enterovirus A89 | EV-A89 | BAN00-10359 | AY697459 |
| *Enterovirus A* | enterovirus A90 | EV-A90 | BAN99-10399 | AY697460 |
| *Enterovirus A* | enterovirus A91 | EV-A91 | BAN00-10406 | AY697461 |
| *Enterovirus A* | enterovirus A92 | EV-A92 | USA/GA99/RJg-7 | EF667344 |
| *Enterovirus A* | enterovirus A114 | EV-A114 | V13-0285/IND/2013 | KU355876 |
| *Enterovirus A* | enterovirus A120 | EV-A120 | MAD-2741-11 (Madagascar) | LK021688 |
| *Enterovirus A* | enterovirus A121 | EV-A121 | V13-0682/IND/2013 | KU355877 |
| *Enterovirus A* | enterovirus A122; simian virus 19 | EV-A122 | M19s | AF326754 |
| *Enterovirus A* | enterovirus A123; simian virus 43 | EV-A123 | OM112t | AF326761 |
| *Enterovirus A* | enterovirus A124; simian virus 46 | EV-A124 | OM22 | AF326764 |
| *Enterovirus A* | enterovirus A125; baboon enterovirus A13 | EV-A125 | A13 | AF326750 |
| *Enterovirus B* | coxsackievirus B3 | CVB3 | Nancy (Connecticut/US/49) | M88483 |
| *Enterovirus B* | coxsackievirus B1 | CVB1 | Japan | M16560 |
| *Enterovirus B* | coxsackievirus B2 | CVB2 | Ohio-1 (Ohio/US/47) | AF081485 |
| *Enterovirus B* | coxsackievirus B4 | CVB4 | JVB (New York/US/51) (Benschoten) | X05690 |
| *Enterovirus B* | coxsackievirus B5 | CVB5 | Faulkner (Kentucky/US/52) | AF114383 |

| | | | | |
|---|---|---|---|---|
| *Enterovirus B* | coxsackievirus B6 | CVB6 | Schmitt (Philippines/53) (1-15-21) | AF039205 |
| *Enterovirus B* | coxsackievirus A9 | CVA9 | Griggs | D00627 |
| *Enterovirus B* | echovirus 1 | E1 | Farouk (Egypt/51) | AF029859 |
| *Enterovirus B* | echovirus 2 | E2 | Cornelis (Connecticut/US/51) | AY302545 |
| *Enterovirus B* | echovirus 3 | E3 | Morrisey (Connecticut/US/51) | AY302553 |
| *Enterovirus B* | echovirus 4 | E4 | Pesacek (Connecticut/US/51) | AY302557 |
| *Enterovirus B* | echovirus 5 | E5 | Noyce (Maine/54) | AF083069 |
| *Enterovirus B* | echovirus 6 | E6 | D'Amori (Rhode Island/55) | AY302558 |
| *Enterovirus B* | echovirus 7 | E7 | Wallace (Ohio) | AY302559 |
| *Enterovirus B* | echovirus 9 | E9 | Hill (Ohio/US/53) | X84981 |
| *Enterovirus B* | echovirus 11 | E11 | Gregory (Ohio) | X80059 |
| *Enterovirus B* | echovirus 12 | E12 | Travis (Philippines/53) | X79047 |
| *Enterovirus B* | echovirus 13 | E13 | Del Carmen (Philippines/53) | AY302539 |
| *Enterovirus B* | echovirus 14 | E14 | Tow (Rhode Island/54) | AY302540 |
| *Enterovirus B* | echovirus 15 | E15 | Ch 96-51 (Charleston) (West Virginia/51) | AY302541 |
| *Enterovirus B* | echovirus 16 | E16 | Harrington (Massachusetts/51) | AY302542 |
| *Enterovirus B* | echovirus 17 | E17 | CHHE-29 (Mexico City) | AY302543 |
| *Enterovirus B* | echovirus 18 | E18 | Metcalf (Ohio) | AF317694 |
| *Enterovirus B* | echovirus 19 | E19 | Burke (Ohio) | AY302544 |
| *Enterovirus B* | echovirus 20 | E20 | JV-1 (Washington DC/55) | AY302546 |
| *Enterovirus B* | echovirus 21 | E21 | Farina (E26D) (Massachusetts/50) | AY302547 |
| *Enterovirus B* | echovirus 24 | E24 | DeCamp (Ohio/56) | AY302548 |
| *Enterovirus B* | echovirus 25 | E25 | JV-4 (Washington DC/57) | AY302549 |
| *Enterovirus B* | echovirus 26 | E26 | Coronel (11-3-6) (Philippines/53) | AY302550 |
| *Enterovirus B* | echovirus 27 | E27 | Bacon (1-36-4) (Philippines/53) | AY302551 |
| *Enterovirus B* | echovirus 29 | E29 | JV-10 (Washington DC/55) | AY302552 |
| *Enterovirus B* | echovirus 30 | E30 | Bastianni (New York/58) | AF162711 |
| *Enterovirus B* | echovirus 31 | E31 | Caldwell (Kansas/55) | AY302554 |
| *Enterovirus B* | echovirus 32 | E32 | PR-10 (Puerto Rico) | AY302555 |
| *Enterovirus B* | echovirus 33 | E33 | Toluca-3 (Mexico/59) | AY302556 |
| *Enterovirus B* | enterovirus B69 | EV-B69 | Toluca-1 (Mexico/59) | AY302560 |
| *Enterovirus B* | enterovirus B73 | EV-B73 | CA55-1988 | AF241359 |
| *Enterovirus B* | enterovirus B74 | EV-B74 | USA/CA75-10213 | AY556057 |
| *Enterovirus B* | enterovirus B75 | EV-B75 | USA/OK85-10362 | AY556070 |

| | | | | |
|---|---|---|---|---|
| *Enterovirus B* | enterovirus B77 | EV-B77 | CF496-99 | AJ493062 |
| *Enterovirus B* | enterovirus B79 | EV-B79 | USA/CA79-10384 | AY843297 |
| *Enterovirus B* | enterovirus B80 | EV-B80 | USA/CA67-10387 | AY843298 |
| *Enterovirus B* | enterovirus B81 | EV-B81 | USA/CA68-10389 | AY843299 |
| *Enterovirus B* | enterovirus B82 | EV-B82 | USA/CA64-10390 | AY843300 |
| *Enterovirus B* | enterovirus B83 | EV-B83 | USA/CA76-10392 | AY843301 |
| *Enterovirus B* | enterovirus B84 | EV-B84 | CIV2003-10603 | DQ902712 |
| *Enterovirus B* | enterovirus B85 | EV-B85 | BAN00-10353 | AY843303 |
| *Enterovirus B* | enterovirus B86 | EV-B86 | BAN00-10354 | AY843304 |
| *Enterovirus B* | enterovirus B87 | EV-B87 | BAN00-10396 | AY843305 |
| *Enterovirus B* | enterovirus B88 | EV-B88 | BAN01-10398 | AY843306 |
| *Enterovirus B* | enterovirus B97 | EV-B97 | BAN99-10355 | AY843307 |
| *Enterovirus B* | enterovirus B98 | EV-B98 | T92-1499 | AB426608 |
| *Enterovirus B* | enterovirus B100 | EV-B100 | BAN2000-10500 | DQ902713 |
| *Enterovirus B* | enterovirus B101 | EV-B101 | CIV03-10361 | AY843308 |
| *Enterovirus B* | enterovirus B106 | EV-B106 | 148/YN/CHN/12 (China/2012) | KF990476 |
| *Enterovirus B* | enterovirus B107 | EV-B107 | TN94-0349 | AB426609 |
| *Enterovirus B* | enterovirus B111 | EV-B111 | Q0011/XZ/CHN/2000 | KF312882 |
| *Enterovirus B* | enterovirus B112 | EV-B112 | GAB130 (chimpanzee/Gabon) | KJ418244 |
| *Enterovirus B* | enterovirus B113 | EV-B113 | GAB653 (mandrill/Gabon) | KJ701249 |
| *Enterovirus B* | enterovirus B114; simian agent 5 | EV-B114 | B165 (vervet monkey) | AF326751 |
| *Enterovirus C* | poliovirus 1 | PV1 | Mahoney | V01149 |
| *Enterovirus C* | poliovirus 1 | PV1 | Sabin (LSc-2ab) | V01150 |
| *Enterovirus C* | poliovirus 2 | PV2 | Lansing (Michigan/37) | M12197 |
| *Enterovirus C* | poliovirus 2 | PV2 | Sabin (P712- Ch-2ab) | X00595 |
| *Enterovirus C* | poliovirus 3 | P-3 | Leon (California/37) | K01392 |
| *Enterovirus C* | poliovirus 3 | PV3 | Sabin (Leon 12a-1-b) | X00925 |
| *Enterovirus C* | coxsackievirus A1 | CVA-1 | T.T. (Tompkins) (Coxsackie/NY/47) | AF499635 |
| *Enterovirus C* | coxsackievirus A11 | CVA-11 | Belgium 1 (Belgium/51) | AF499636 |
| *Enterovirus C* | coxsackievirus A13 | CVA-13 | Flores (Mexico/52) | AF499637 |
| *Enterovirus C* | coxsackievirus A13; coxsackievirus A18 | CVA-13 | G-13 (South Africa/50) | AF499640 |
| *Enterovirus C* | coxsackievirus A17 | CVA-17 | G12 (South Africa/51 | AF499639 |
| *Enterovirus C* | coxsackievirus A19 | CVA-19 | NIH-8663 (Dohi) (Japan/52) | AF499641 |
| *Enterovirus C* | coxsackievirus A20 | CVA-20 | IH-35 (New York/55) | AF499642 |

| | | | | |
|---|---|---|---|---|
| *Enterovirus C* | coxsackievirus A21 | CVA-21 | Kuykendall (California/52) | AF546702 |
| *Enterovirus C* | coxsackievirus A22 | CVA-22 | Chulman (New York/55) | AF499643 |
| *Enterovirus C* | coxsackievirus A24 | CVA-24 | Joseph | EF026081 |
| *Enterovirus C* | enterovirus C96 | EV-C96 | BAN00-10499 | EF015886 |
| *Enterovirus C* | enterovirus C99 | EV-C99 | USA-GA84-10636 | EF555644 |
| *Enterovirus C* | enterovirus C102 | EV-C102 | BAN99-10424 | EF555645 |
| *Enterovirus C* | enterovirus C105 | EV-C105 | PER153 (Peru/2010) | JX393302 |
| *Enterovirus C* | enterovirus C109 | EV-C109 | NICA08-4327 | GQ865517 |
| *Enterovirus C* | enterovirus C113 | EV-C113 | BBD-48/Bangladesh/2009 | KC344833 |
| *Enterovirus C* | enterovirus C116 | EV-C116 | 126/Russia/2010 | JX514942 |
| *Enterovirus C* | enterovirus C117 | EV-C117 | LIT22/Vilnius (Lithuania/2011) | JX262382 |
| *Enterovirus C* | enterovirus C118 | EV-C118 | ISR10 (Israel/2011) | JX961708 |
| *Enterovirus D* | enterovirus D68 | EV-D68 | Fermon | AY426531 |
| *Enterovirus D* | enterovirus D70 | EV-D70 | J670/71 (Japan/71) | D00820 |
| *Enterovirus D* | enterovirus D94 | EV-D94 | E210 (Egypt) | DQ916376 |

Table S4 Phylogenetical classification of the previously tested CVB5 variants[2] by VP1, 5’UTR, non-structural proteins, and 3’UTR regions. The variants were classified into two genogroups A and B according to the bootstrapped and neighbor-joining phylogenetic trees with 1000 replicates constructed by MEGA X software[3]. Highlighted letters indicate that the classification of a certain variant is as same as the VP1-based classification.

| **Variant** | **VP1[a]** | **5’UTR** | **Non-structural proteins** | **3’UTR** |
|---|---|---|---|---|
| CVB5.1 | A | A | B | B |
| CVB5.2 | A | A | B | B |
| CVB5.3 | B | B | A | A |
| CVB5.4 | A | A | A | A |
| CVB5.5 | A | A | A | A |
| CVB5.6 | B | B | B | B |
| CVB5.7 | B | B | B | A |
| CVB5.8 | B | B | B | A |
| CVB5.9 | A | A | A | A |
| CVB5.10 | B | B | B | B |
| CVB5.11 | A | A | B | B |
| CVB5.12 | B | B | B | B |
| CVB5.Faulkner | A | B | A | A |

[a] The results of classification is according to our previous studies [2].

Table S5 Primers used for site-directed mutagenesis.

| Name | Sequence (5' - 3') |
|---|---|
| VP2.L137I-fd | GGG TGC GCG ACG ATA GCA AAC AAA CCT GA |
| VP2.L137I-rv | TCA GGT TTG TTT GCT ATC GTC GCG CAC CC |
| VP2.E156D-fd | CGC CAA CGT GTT TGA CTC TCA GAA CTC TTC |
| VP2.E156D-rv | GAA GAG TTC TGA GAG TCA AAC ACG TTG GCG |
| VP2.S160T-fd | TTG AGT CTC AGA ACA CGT CGG GAC AAA CAG |
| VP2.S160T-rv | CTG TTT GTC CCG ACG TGT TCT GAG ACT CAA |
| VP2.K260R-fd | CGC TTG GCT GGC AGA CAG GGG CTT CCC ACA |
| VP2.K260R-rv | TGT GGG AAG CCC CTG TCT GCC AGC CAA GCG |
| VP3.M63L-fd | CTG AGG GGA AGG TGC TCT CCA TTG AAG CCT A |
| VP3.M63L-rv | TAG GCT TCA ATG GAG AGC ACC TTC CCC TCA G |
| VP3.T88I-fd | GGA TTC CCA CTG ATC CCA GGG GCT AGT AG |
| VP3.T88I-rv | CTA CTA GCC CCT GGG ATC AGT GGG AAT CC |
| VP1.V156I-fd | CGT GCC CAC AAA AAT CAA CAG CTA CAG TTG |
| VP1.V156I-rv | CAA CTG TAG CTG TTG ATT TTT GTG GGC ACG |
| VP1.M180I-fd | TGC CCC ACC TCG GAT CTC GAT ACC CTT CAT TAG |
| VP1.M180I-rv | CTA ATG AAG GGT ATC GAG ATC CGA GGT GGG GCA |
| VP1.D276E-fd | GAG GGC AGA ACA GAG ATA ACG ACC ATG C |
| VP1.D276E-rv | GCA TGG TCG TTA TCT CTG TTC TGC CCT C |
| VP1.T279A-fd | GAA CAG ACA TAA CGG CTA TGC AAA CCA CTG |
| VP1.T279A-rv | CAG TGG TTT GCA TAG CCG TTA TGT CTG TTC |

Table S6 Confirmation of the introduced mutation by sanger sequencing.

| **Mutant** | **Position of respective amino acid residue (nt)** | **Genome sequence of corresponding residue** | |
|---|---|---|---|
| | | **CVB5F.cas** | **Mutant** |
| CVB5F.cas.VP2.L137I | 1358-1360 | CTG | ATA |
| CVB5F.cas.VP2.E156D | 1415-1717 | GAG | GAC |
| CVB5F.cas.VP2.S160T | 1427-1429 | TCT | ACG |
| CVB5F.cas.VP2.K260R | 1727-1729 | AAA | AGA |
| CVB5F.cas.VP3.M63L | 1919-1921 | ATG | CTC |
| CVB5F.cas.VP3.T88I | 1994-1996 | ACT | ATC |
| CVB5F.cas.VP1.V156I | 2912-2914 | GTA | ATY[a] |
| CVB5F.cas.VP1.M180I | 2984-2986 | ATG | ATC |
| CVB5F.cas.VP1.D276E | 3272-3274 | GAC | GAG |
| CVB5F.cas.VP1.T279A | 3281-3283 | ACC | GCT |
| CVB5F.cas.genogroupB [b] | - | - | - |

[a] Y represents pyrimidine (C or T). Both ATC and ATT codes isoleucine(I).

[b] All the substitution was successfully introduced.

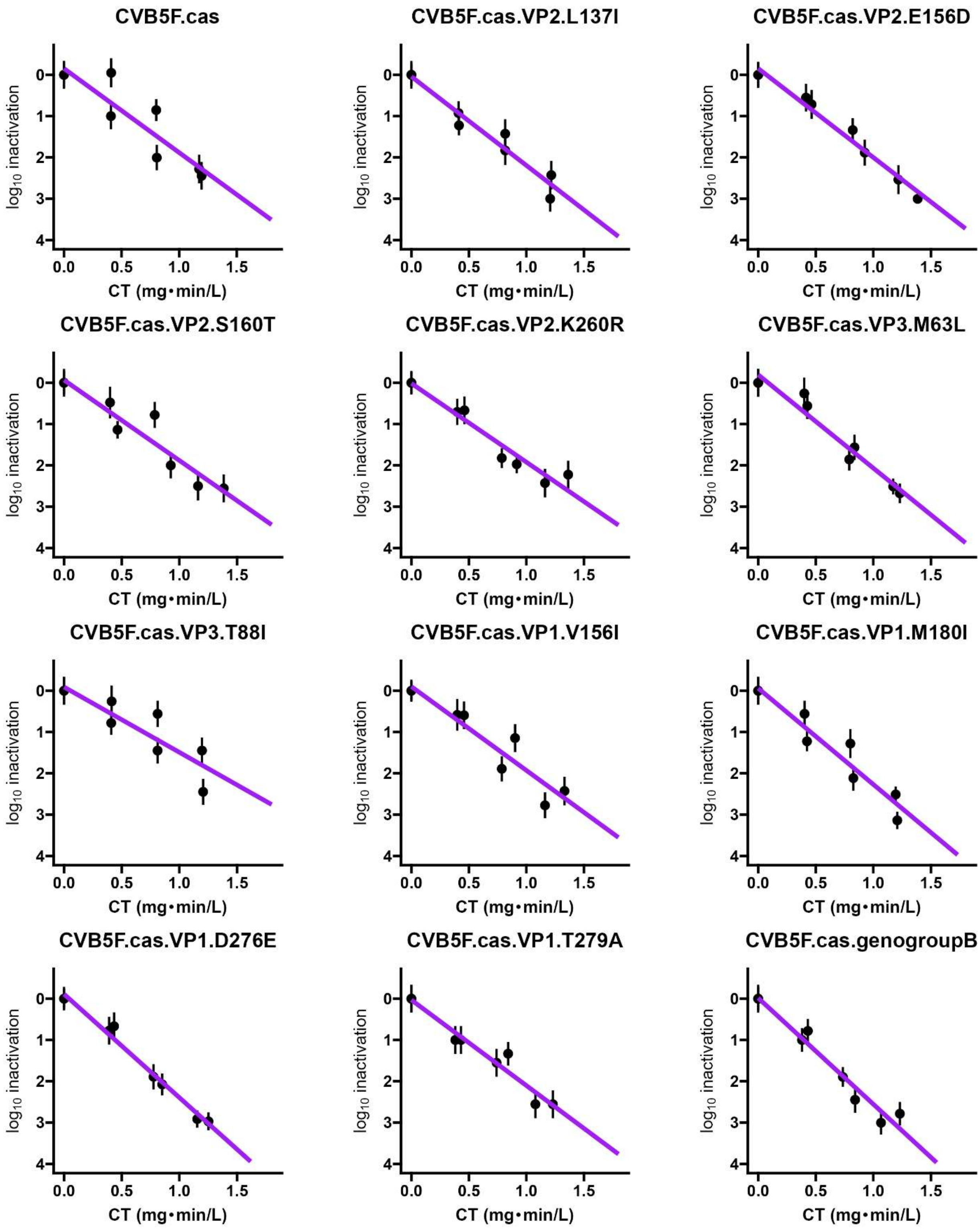


Figure S1 Inactivation curves for free chlorine. The experiments were performed at pH 7.0 at 20°C. The purple line indicates the regression line. The log inactivation was plotted versus CT values (mg min $L^{-1}$) along with error bars representing theoretical standard errors of MPN values.

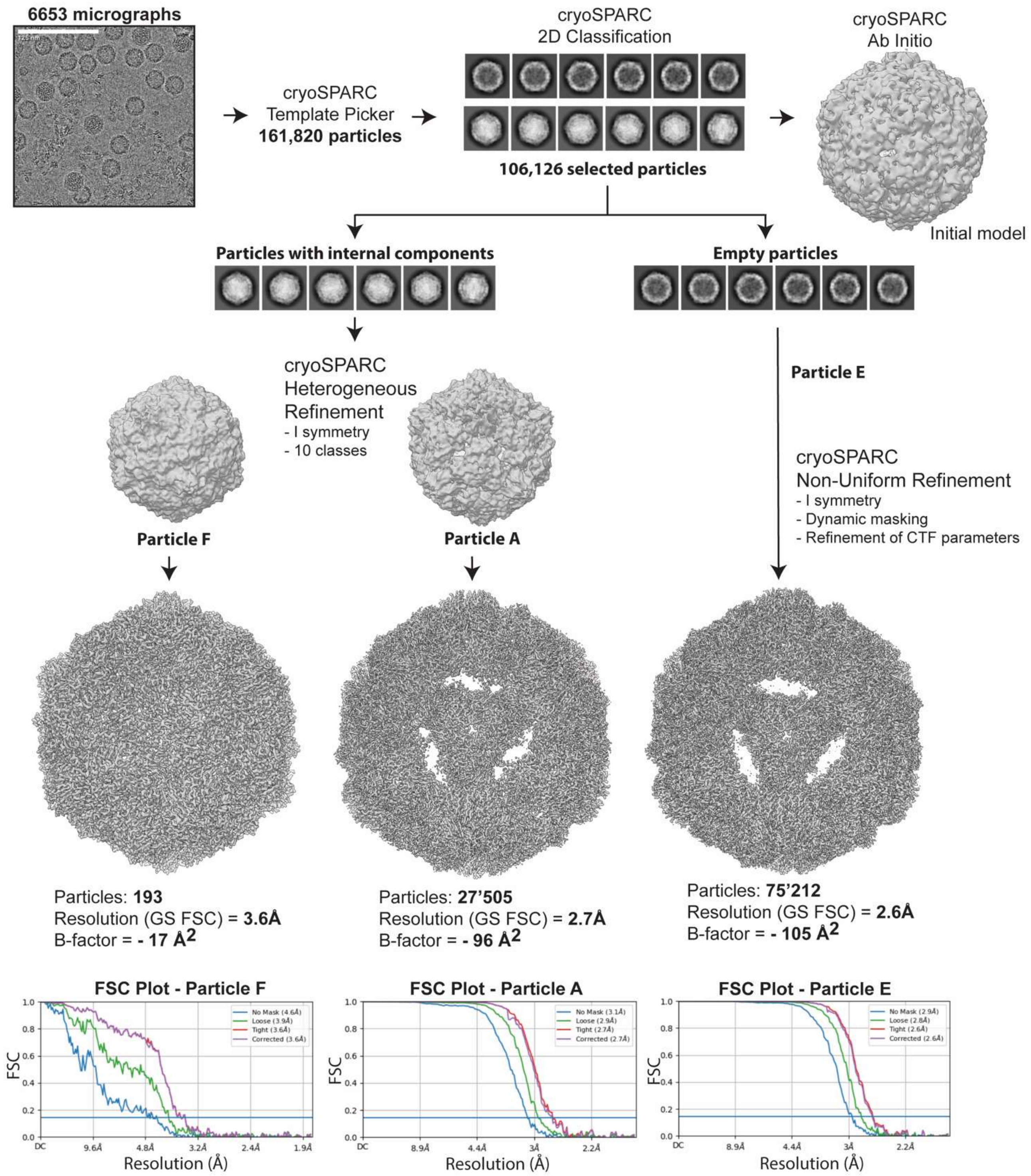


Figure S2 The workflow used to process cryoEM data.

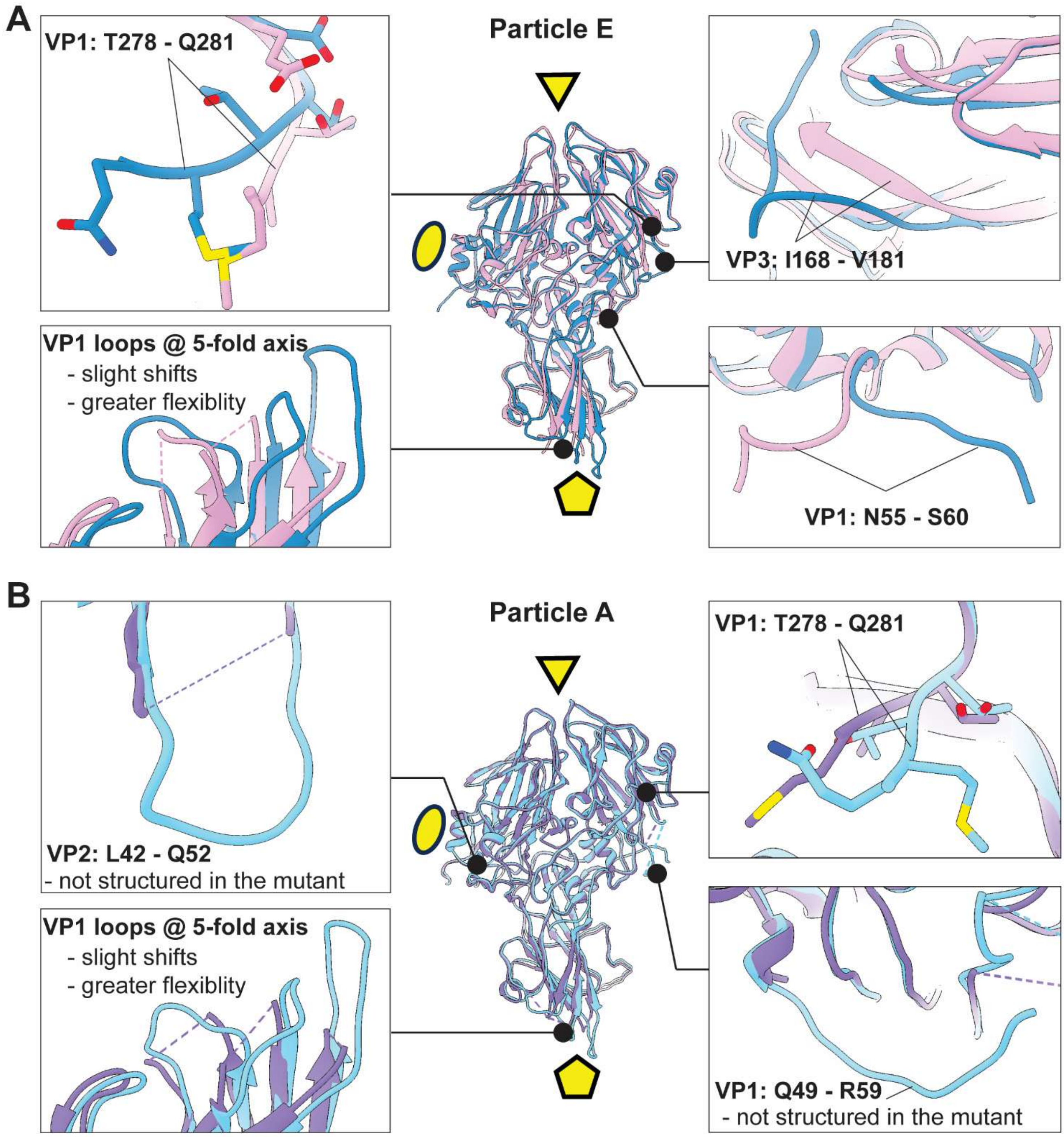


Figure S3 (A) Comparison of the E states in CVB5F.cas.genogroupB (plum) to the previously published structures of the E state CVB5 (PDB IDs: 7WL3; blue). (B) Comparison of the A states in CVB5F.cas.genogroupB (purple) to the previously published structures of the A state CVB5 (PDB IDs: 7XB2; light blue). In both panels, the overlay of the two structures is shown in the middle, and the close-up views of the most notable differences are on each side. For simplicity, only 1 protomer is shown with the locations of 2-, 3-, and 5-fold symmetry axis indicated with ellipse, triangle and pentamer, respectively.

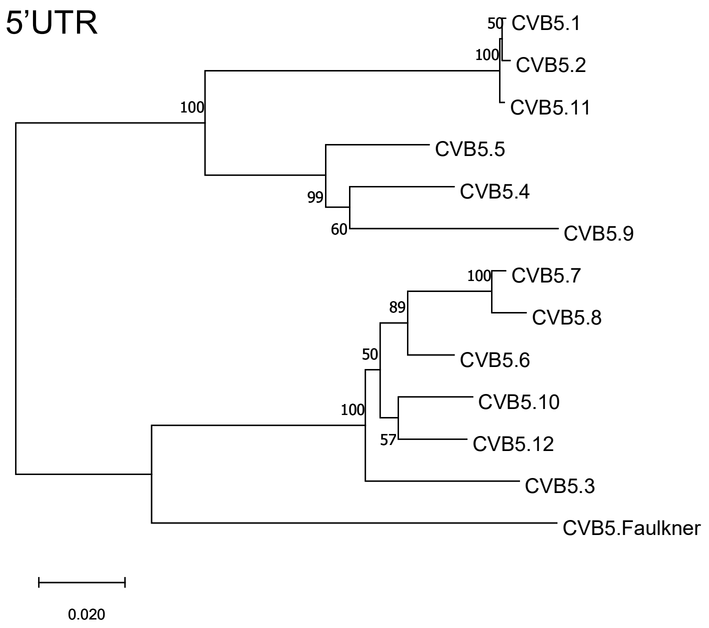


Figure S4 Phylogenetic tree constructed for 5’UTR region of CVB5 variants. The percentages of the replicate trees in which the associated taxa clustered together in the bootstrap test (1000 replicates) are shown next to the branches.
